# ER Stress-Response Signaling Regulates Chamber-Specific Growth between Right and Left Ventricles during Postnatal Development

**DOI:** 10.64898/2026.07.15.738823

**Authors:** Boyang Zhang, Michal B. Juda, Jijun Huang, Douglas Chapski, Adrian Arrieta, Isaac Rodney, Jinge Liu, Michael Leone, Tzung Hsiai, Yibin Wang, Tomohiro Yokota

**Author notes:** Correspondence Tomohiro Yokota, PhD MRL 1619, 675 Charles E Young Drive S, University of California, Los Angeles, CA 90095. Lead Contact: Further information and requests for resources and reagents should be directed to and will be fulfilled by Tomohiro Yokota.

## Abstract

**Background:** Differential growth between the left (LV) and right ventricles (RV) is a cornerstone of normal heart morphogenesis after birth, leading to the relatively larger and dominant LV over RV in the adult heart regarding size and function. Yet, little is known about the factors that regulate this chamber-specific growth.

**Methods:** We used both loss-and gain-of-function mouse models, achieved through genetic or pharmacological manipulation of IRE1α or Xbp1 in cardiomyocytes. We also used primary cultured neonatal cardiomyocytes to explore the roles of IRE1α, spliced Xbp1 (sXbp1: activated form), and newly identified sXbp1 downstream targets. In addition, we generated heart-specific mosaic mutant mouse models using CRISPR/Cas9/AAV9-based somatic mutagenesis to elucidate the roles of sXbp1 downstream targets in cardiomyocytes.

**Results:** Pharmacological inactivation of IRE1α and genetic depletion of Xbp1 resulted in a smaller LV size, due to decreased cardiomyocyte proliferation and hypertrophic growth, as well as increased cardiomyocyte death. These effects were not observed in the RV. Cardiomyocyte-specific induction of IRE1α or sXbp1 led to increased ventricular size in both ventricles, through enhanced cardiomyocyte proliferation and hypertrophic growth in both LV and RV, and reduced apoptosis in the RV. We identified two ER resident transmembrane proteins, Vimp and Rpn2, as direct binding partners of sXbp1 in targeted gene regulation at the chromatin level. CRISPR/Cas9/AAV9-based somatic mutagenesis mouse models for Vimp and Rpn2 revealed that both genes regulate cardiomyocyte proliferation, hypertrophic growth, and death. We also observed accumulated misfolded proteins in these two mutant hearts.

**Conclusions**

We demonstrate that the IRE1α-Xbp1-Vimp/Rpn2 axis regulates differential ventricular size between LV and RV during postnatal development by orchestrating cardiomyocyte proliferation, hypertrophic growth, and death through regulating protein homeostasis.

**Clinical Perspective:** *What Is New:*

- IRE1α-Xbp1 axis is dominantly activated in the LV cardiomyocyte during the postnatal period in mouse heart.
- IRE1α-Xbp1 mediated ER stress signaling increases cardiomyocyte proliferation and hypertrophic growth and decreases apoptosis in the postnatal period.
- Activated Xbp1 directly regulates LV-specific cardiomyocyte protein homeostasis via interaction with ER membrane targeted Vimp and Rpn2.

*What Are the Clinical Implications?:*

- Differential heart growth patterns between the LV and RV are critical for normal morphogenesis and function of each ventricle.
- Control of protein homeostasis by modulating ER stress signaling could be a potential therapeutic approach for single-chamber heart diseases.

## Introduction

During fetal development, the right ventricle (RV) is comparable in size to, or even exceeds, the left ventricle (LV) in cardiac output, as both chambers function within a common circulation^1,2^. After birth, this balance shifts rapidly: the LV surpasses the RV in contractile function, chamber size, and wall thickness^3–6^. This dramatic postnatal remodeling accommodates the abrupt separation of systemic and pulmonary circulations, imposing high peripheral resistance on the LV while pulmonary resistance declines for the RV ^7,8^. The resulting differential mechanical and hemodynamic load drives chamber-specific remodeling through coordinated cardiomyocyte proliferation, hypertrophy, and apoptosis, resulting in a markedly larger LV in adolescent and adult hearts ^4,9^. Although this asymmetric growth is a hallmark of postnatal heart development, the underlying molecular mechanisms remain poorly defined. Current models emphasize extrinsic cues, including intracavitary pressure, shear stress, hormonal signaling, vascular growth, and sympathetic innervation, as drivers of chamber-specific cardiomyocyte proliferation, hypertrophy, and apoptosis ^9–12^. Yet, the intracellular networks that confer LV-versus RV-specific responses remain unknown.

The endoplasmic reticulum (ER) is an essential organelle responsible for protein folding, lipid synthesis, and calcium homeostasis. Disruption of ER function by conditions such as hypoxia, oxidative stress, nutrient deprivation, or increased protein synthesis leads to the accumulation of misfolded proteins, causing ER stress. To restore cellular homeostasis, cells activate the unfolded protein response (UPR), a signaling network mediated by three major ER stress sensors: PERK, IRE1α, and ATF6. The outcome of these pathways will reduce protein overload, enhance protein-folding capacity, and promote degradation of damaged proteins^13^. Importantly, ER stress signaling plays an important role in regulating cell proliferation, growth, and survival in a context specific manner^14–16^. Moderate activation of the UPR supports cellular adaptation, survival, and biosynthetic activity required for proliferation, whereas prolonged or excessive ER stress can trigger growth arrest and apoptosis^17^. UPR signaling also interacts with metabolic and growth-regulatory pathways, including mTOR and autophagy, linking ER homeostasis to cellular growth control ^18^. In addition to its role in stress adaptation, ER stress signaling contributes to tissue development and organ homeostasis ^19–21^. In the heart, controlled UPR activation is important for cardiomyocyte survival, protein quality control, and adaptation to developmental and physiological stress^22^. The IRE1α-XBP1 signaling pathway has emerged as a key regulator of cardiac homeostasis by promoting adaptive gene expression involved in protein folding, secretion, metabolism, and angiogenesis ^23^. XBP1 has also been implicated in cardiomyocyte growth and protection during ischemic and hypertrophic stress, while dysregulated or chronic ER stress contributes to cardiac hypertrophy, fibrosis, and heart failure ^24–29^.

Here, we demonstrate that IRE1α-Xbp1 pathway regulates chamber-specific growth between LV and RV. Animals lacking IRE1α activity or Xbp1 in cardiomyocytes exhibit a significant reduction in LV to comparable RV size. On the other hand, animals with cardiomyocyte specific expression of IRE1α or spliced Xbp1, the active form of Xbp1, show a significant enlargement of both ventricles, leading to the loss of RV specific features. These impact is at least in part associated with two newly identified Xbp1 binding proteins, Vimp and Rpn2 on ER membrane. These findings establish IRE1α-Xbp1 axis in ER stress signaling as an intrinsic mechanism for the physiological repgoramming and chamber-specific morphogenesis during postnatal stages. These insights might have implications for the treatment of single-chamber congenital heart diseases.

## Results

### IRE1α-sXbp1 axis regulates cardiomyocyte proliferation, hypertrophic growth, and cell death in cultured cardiomyocytes

In our previous study, IRE1α is identifed as a potential downstream regulator of p38 MAPK in the regulation of chamber-specific postnatal heart growth^30^. In mouse LV and RV tissues obtained from postnatal day 1 (P1) to P28, we measured the expression of spliced Xbp1 (sXbp1), a downstream product of activated IRE1α. Tissue levels of sXbp1 were higher in the LV than in the RV at P1 (**Figure 1A)** and remained significantly elevated in isolated LV myocytes than RV at P3 postnatal time point (**Figure 1B**). To confirm these results, we performed RNA-fluorescence in situ hybridization (RNA-FISH) analysis with immunofluorescence probe for cardiac troponin I as a cardiomyocyte marker ^31^ to determine the specific expression of sXbp1 in cardiomyocytes. The results showed robust enrichment of Xbp1 in cardiomyocyte nuclei in the LV vs. the RV, indicating chamber specific induction of spliced and activated product of sXbp1 (**Figure 1C**). This LV-specific expression pattern of sXbp1 implicates its role in chamber-specific growth within postnatal hearts.

**Figure 1.**
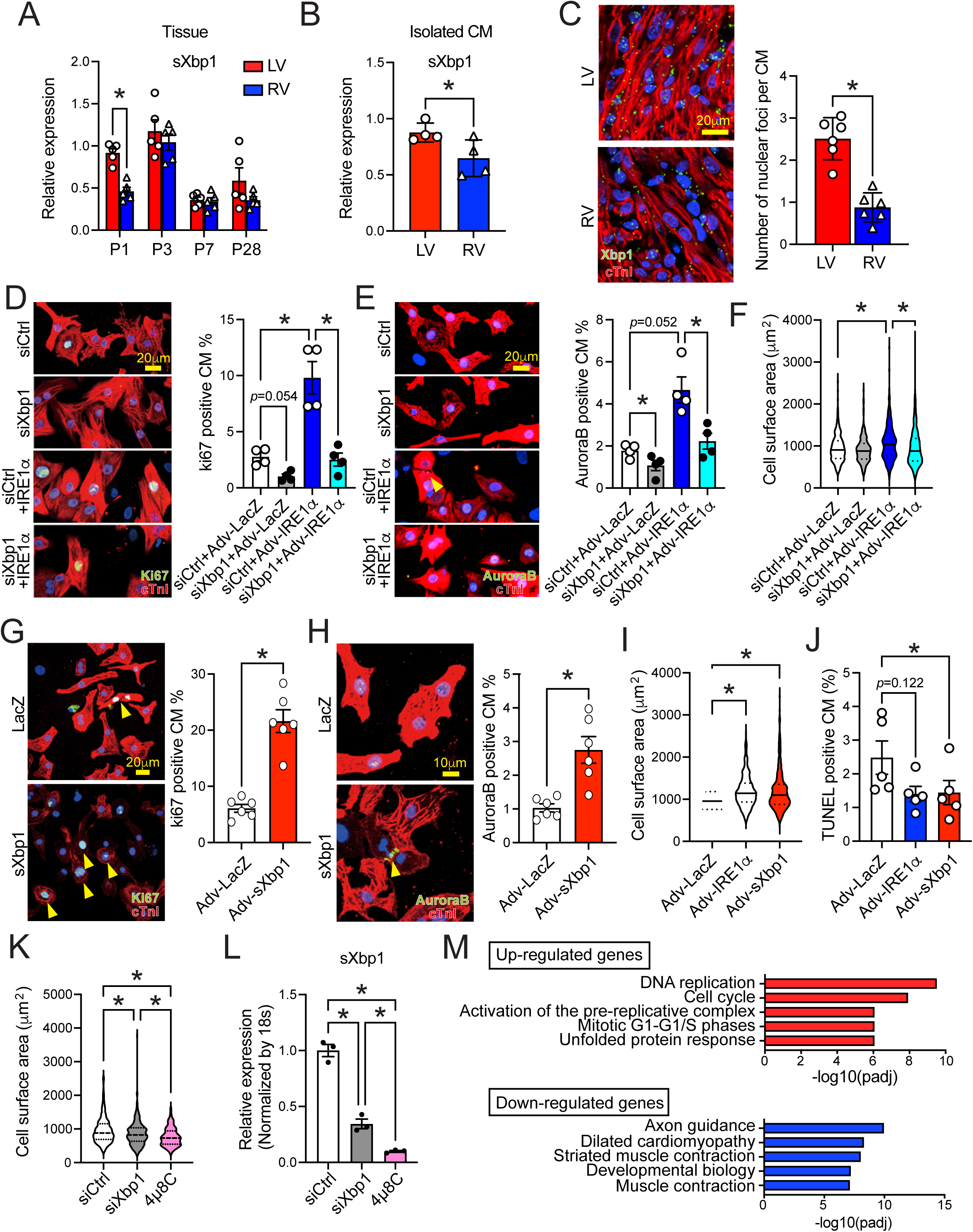
Temporal expression of spliced Xbp1 during the postnatal stage and its role in neonatal cardiomyocytes. **(A)** Expression of spliced Xbp1 (sXbp1) in LV and RV at postnatal day 1 (P1), P3, P7, and P28 (n=5). **(B)** Expression of sXbp1 in isolated cardiomyocytes from LV and RV at P3 (n=4). **(C)** RNA-FISH to demonstrate expression of Xbp1 in the ventricles at P3. Quantification of Xbp1 signal in nuclei in each cardiomyocyte (n=6). **(D,E)** Immunocytochemistry against Ki67 and Aurora B **(E)** on cardiomyocytes treated with adenovirus (adv)-IRE1α and siRNA against Xbp1 (n=4). **(F)** Cell surface area of cardiomyocytes treated with adv-IRE1α and siRNA against Xbp1 (n=4, 100 cardiomyocytes per each experiment). **(G,H)** Immunocytochemistry against ki67 and Aurora B **(H)** on cardiomyocytes treated with adv-sXbp1 (n=6). **(I)** Cell surface area and cell death ratio **(J)** of cardiomyocytes treated with adv-IRE1a and adv-sXbp1 (n=4 and n=5, respectively). **(K)** Cell surface area and sXbp1 expression **(L)** of cardiomyocytes treated with siRNA against Xbp1 and an IRE1a inhibitor, 4µ8C (n=3). **(M)** Pathway analysis on cardiomyocytes treated with adv-sXbp1 (n=3). All data are shown as mean ± SEM, *p<0.05

To demonstrate that Xbp1 is necessary for IRE1α-induced proliferation in cultured cardiomyocytes, we silenced Xbp1 in neonatal rat ventricle cardiomyocytes by transfecting siRNA against Xbp1 in the presence of IRE1α overexpression. We confirmed that Xbp1 silencing blunted basal and IRE1α-induced proliferation in neonatal myocytes(**Figure 1D and 1E**). We also demonstrated that the silencing of Xbp1 blocked the cardiomyocyte hypertrophy, a hallmark of neonatal myocyte maturation, induced by IRE1α expression (**Figure 1F**). Therefore, higher activities in IRE1α-Xbp1 regulatory axis in LV promote chamber specific myocyte proliferation and hypertrophic growth during postnatal heart development.

To demonstrate that sXbp1 is sufficient to induce proliferation and hypertrophic growth, we treated cardiomyocytes with adenovirus expressing sXbp1 (Adv-sXbp1) ^24^. We observed that sXbp1 expression induced both proliferation and hypertrophic growth in cultured neonatal cardiomyocytes (**Figure 1G and 1H**). Gene expression profile revealed that sXbp1-expression was associated with enhanced cell proliferation and reduced cardiac maturation markers in a dose-dependent manner (**Figure S1A**). This effect was cardiomyocyte specfici as changes in proliferation was observed in neonatal ventricular fibroblasts upon sXbp1 expression (**Figure S1B-D**). In addition to cell proliferation, chamber-specific growth is contributed by programmed cell death as well^9,30^, we performed TUNEL staining on neonatal cardiomyocytes treated with Adv-IRE1α or Adv-sXbp1 and found that their expression reduced apoptosis levels (**Figure 1J**). Furthermore, to demonstrate the effect of IRE1α inactivation or Xbp1 silencing on hypertrophic growth, we treated neonatal cardiomyocytes with siRNA against Xbp1 or 4µ8C ^32^, an IRE1α inhibitor, and found both treatments reduced cardiomyocyte size (**Figure 1K**). Interestingly, the total levels of sXbp1 were inversely correlated with cardiomyocyte size treated with Xbp1 silencing or 4µ8C (**Figure 1L**). To further demonstrate the role of sXbp1 in an unbiased manner, we performed bulk RNA-seq on sXbp1-expressing cardiomyocytes. 1,196 up-regulated genes and 1,259 down-regulated genes were detected. From these differentially expressed genes, cellular pathways associated with proliferation and ER-related protein homeostasis, including genes involved in the unfolded protein response (UPR), were upregulated, while maturation and sarcomeric structure genes were downregulated by sXbp1 induction (**Figure 1M**, **Table 1, and Supplemental Table 1**). Taken together, Xbp1 is necessary for IRE1α-induced myocyte proliferation and hypertrophic growth, and IRE1α-sXbp1 axis is a potent regulator for neonatal myocyte proliferation, hypertrophy, and programmed cell death. These results suggest that differential growth between LV and RV during the postnatal period likely involves ER-regulated protein homeostasis.

**Table 1:**
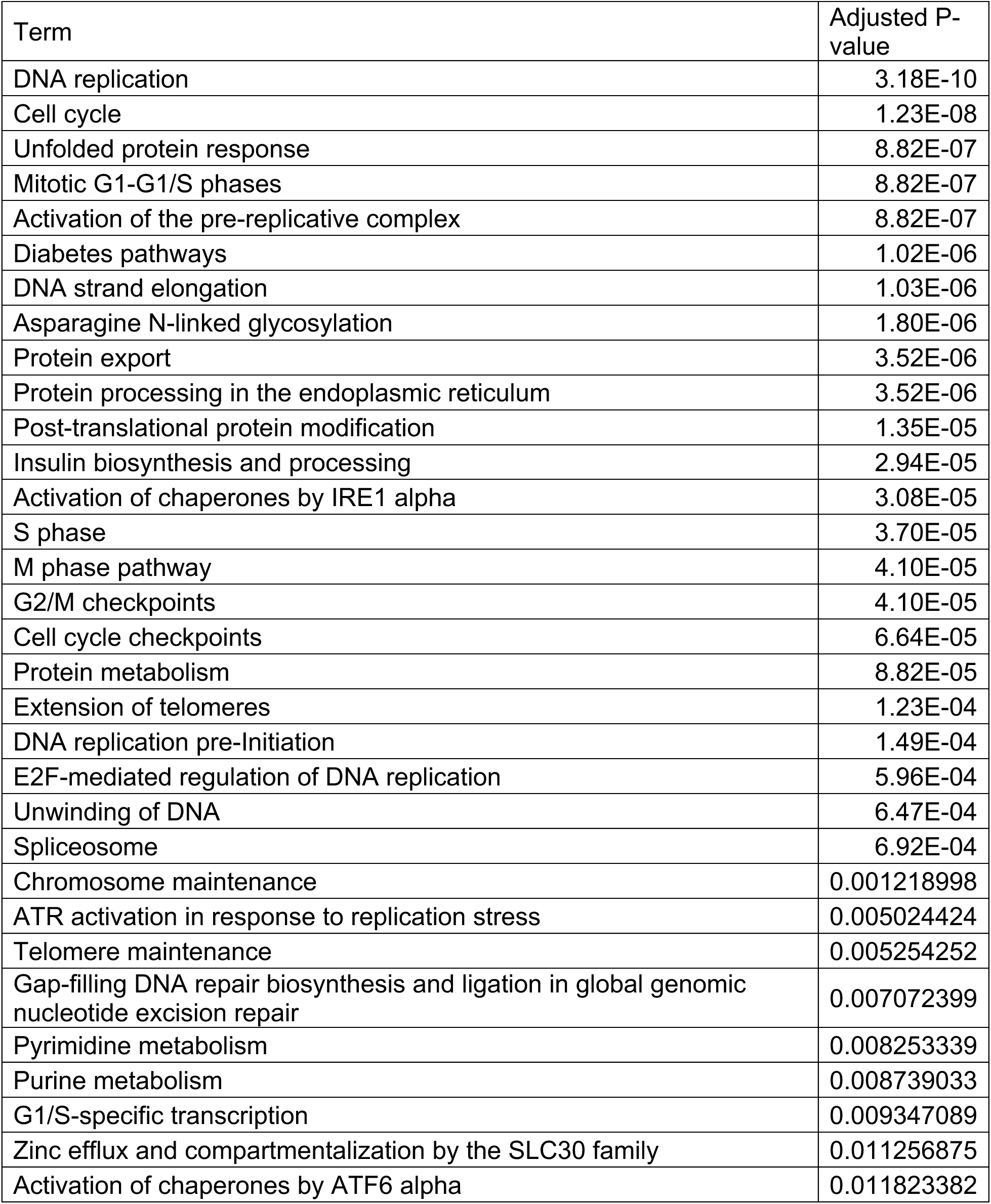
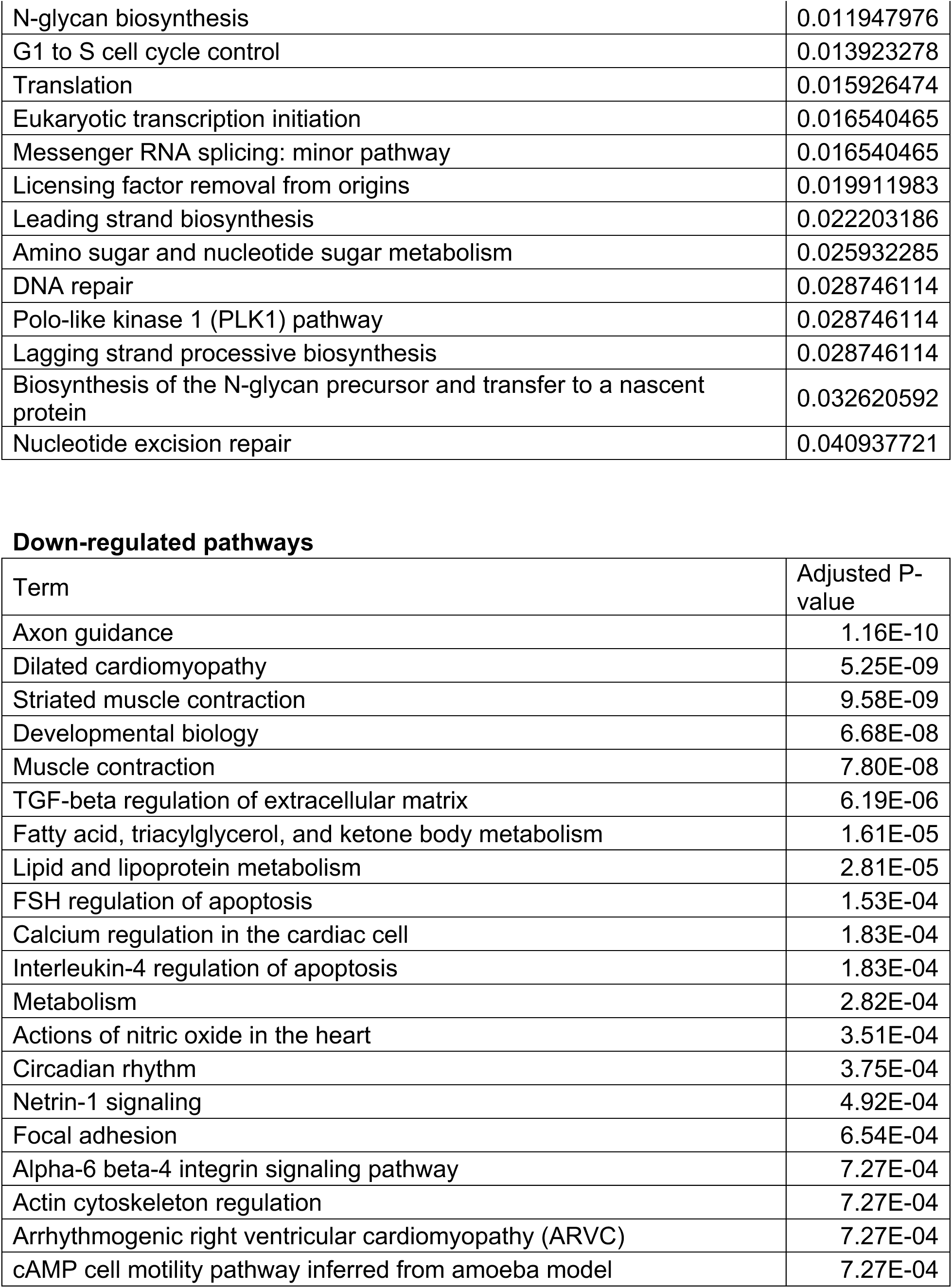
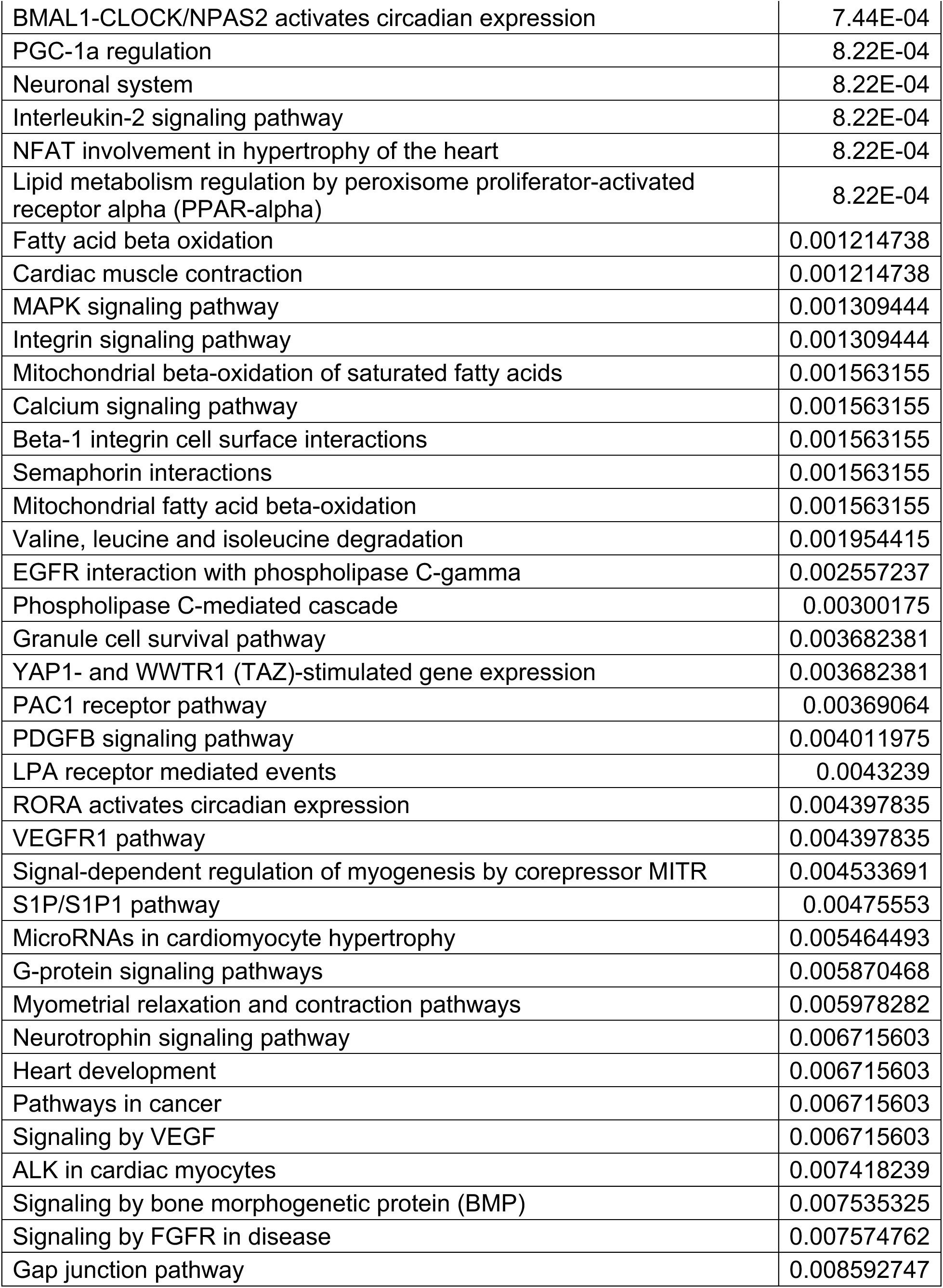
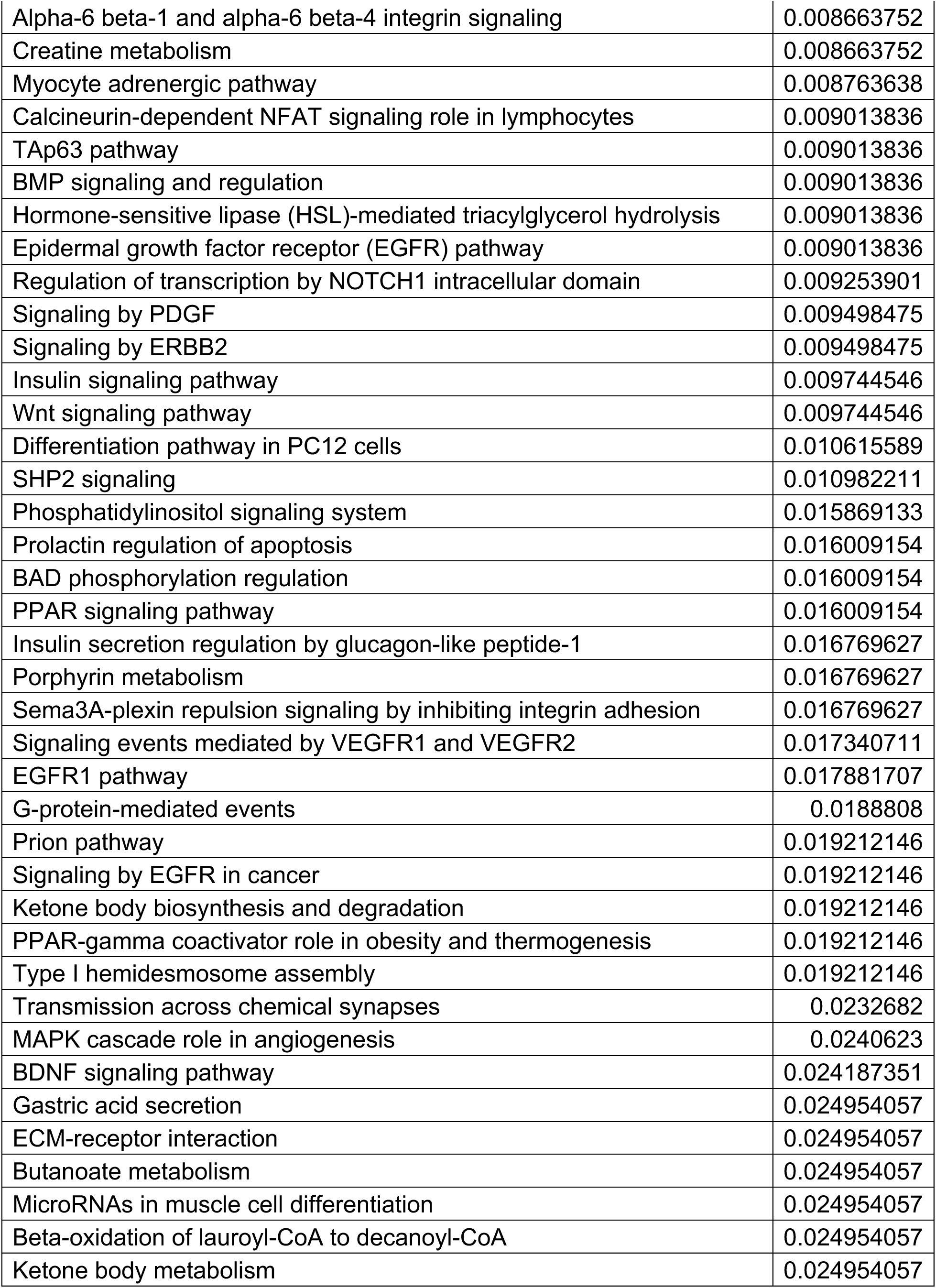
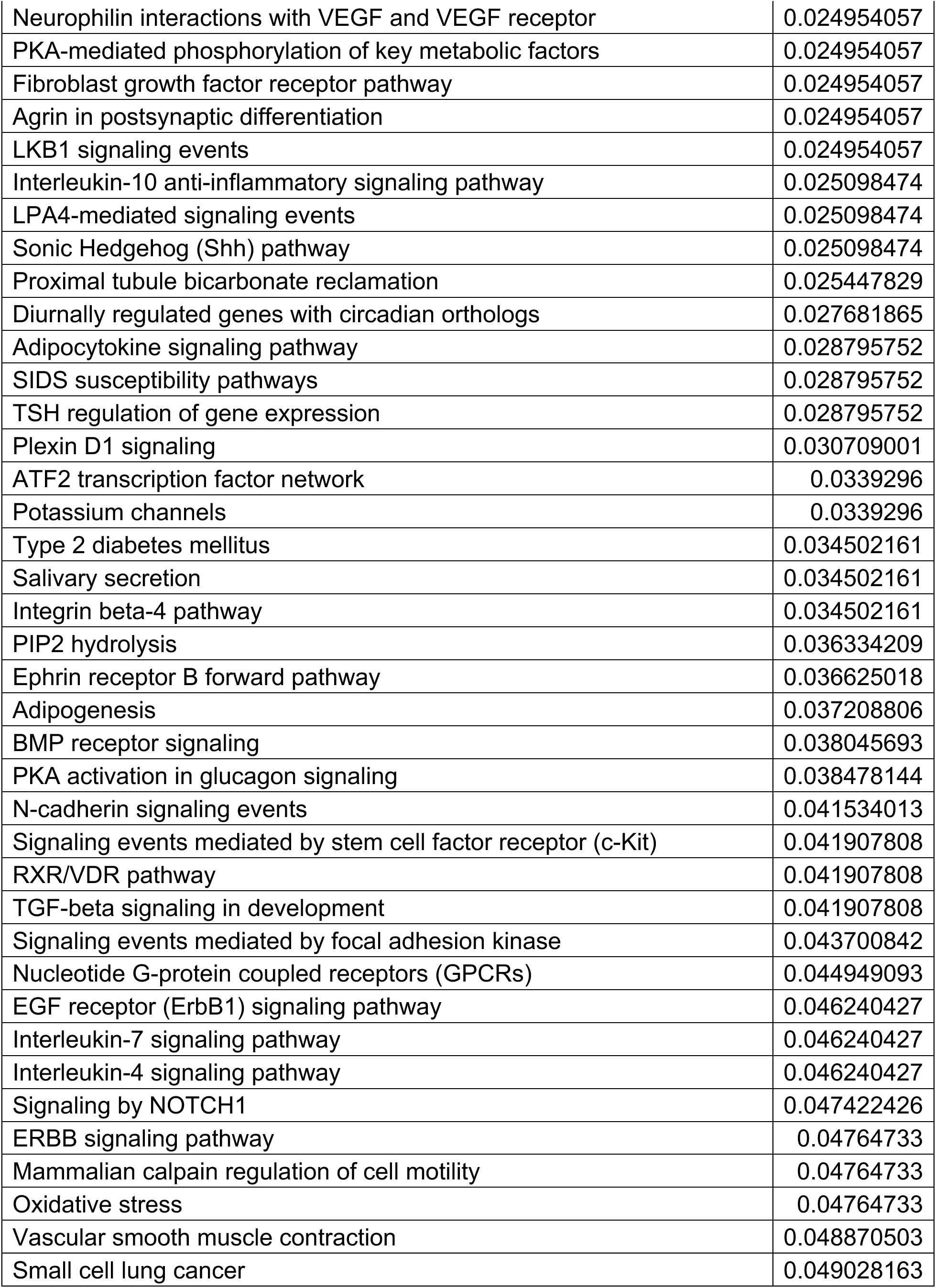
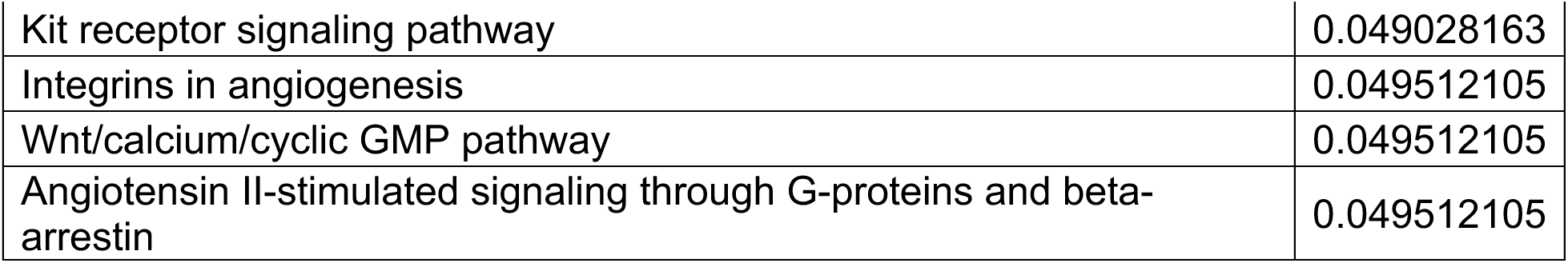
List of pathway analysis from sXbp1-induced neonatal cardiomyocytes Up-regulated pathways.

| Term | Adjusted P-value |
| --- | --- |
| DNA replication | 3.18E-10 |
| Cell cycle | 1.23E-08 |
| Unfolded protein response | 8.82E-07 |
| Mitotic G1-G1/S phases | 8.82E-07 |
| Activation of the pre-replicative complex | 8.82E-07 |
| Diabetes pathways | 1.02E-06 |
| DNA strand elongation | 1.03E-06 |
| Asparagine N-linked glycosylation | 1.80E-06 |
| Protein export | 3.52E-06 |
| Protein processing in the endoplasmic reticulum | 3.52E-06 |
| Post-translational protein modification | 1.35E-05 |
| Insulin biosynthesis and processing | 2.94E-05 |
| Activation of chaperones by IRE1 alpha | 3.08E-05 |
| S phase | 3.70E-05 |
| M phase pathway | 4.10E-05 |
| G2/M checkpoints | 4.10E-05 |
| Cell cycle checkpoints | 6.64E-05 |
| Protein metabolism | 8.82E-05 |
| Extension of telomeres | 1.23E-04 |
| DNA replication pre-Initiation | 1.49E-04 |
| E2F-mediated regulation of DNA replication | 5.96E-04 |
| Unwinding of DNA | 6.47E-04 |
| Spliceosome | 6.92E-04 |
| Chromosome maintenance | 0.001218998 |
| ATR activation in response to replication stress | 0.005024424 |
| Telomere maintenance | 0.005254252 |
| Gap-filling DNA repair biosynthesis and ligation in global genomic nucleotide excision repair | 0.007072399 |
| Pyrimidine metabolism | 0.008253339 |
| Purine metabolism | 0.008739033 |
| G1/S-specific transcription | 0.009347089 |
| Zinc efflux and compartmentalization by the SLC30 family | 0.011256875 |
| Activation of chaperones by ATF6 alpha | 0.011823382 |
| N-glycan biosynthesis | 0.011947976 |
| G1 to S cell cycle control | 0.013923278 |
| Translation | 0.015926474 |
| Eukaryotic transcription initiation | 0.016540465 |
| Messenger RNA splicing: minor pathway | 0.016540465 |
| Licensing factor removal from origins | 0.019911983 |
| Leading strand biosynthesis | 0.022203186 |
| Amino sugar and nucleotide sugar metabolism | 0.025932285 |
| DNA repair | 0.028746114 |
| Polo-like kinase 1 (PLK1) pathway | 0.028746114 |
| Lagging strand processive biosynthesis | 0.028746114 |
| Biosynthesis of the N-glycan precursor and transfer to a nascent protein | 0.032620592 |
| Nucleotide excision repair | 0.040937721 |

Down-regulated pathways
| Term | Adjusted P-value |
| --- | --- |
| Axon guidance | 1.16E-10 |
| Dilated cardiomyopathy | 5.25E-09 |
| Striated muscle contraction | 9.58E-09 |
| Developmental biology | 6.68E-08 |
| Muscle contraction | 7.80E-08 |
| TGF-beta regulation of extracellular matrix | 6.19E-06 |
| Fatty acid, triacylglycerol, and ketone body metabolism | 1.61E-05 |
| Lipid and lipoprotein metabolism | 2.81E-05 |
| FSH regulation of apoptosis | 1.53E-04 |
| Calcium regulation in the cardiac cell | 1.83E-04 |
| Interleukin-4 regulation of apoptosis | 1.83E-04 |
| Metabolism | 2.82E-04 |
| Actions of nitric oxide in the heart | 3.51E-04 |
| Circadian rhythm | 3.75E-04 |
| Netrin-1 signaling | 4.92E-04 |
| Focal adhesion | 6.54E-04 |
| Alpha-6 beta-4 integrin signaling pathway | 7.27E-04 |
| Actin cytoskeleton regulation | 7.27E-04 |
| Arrhythmogenic right ventricular cardiomyopathy (ARVC) | 7.27E-04 |
| cAMP cell motility pathway inferred from amoeba model | 7.27E-04 |
| BMAL1-CLOCK/NPAS2 activates circadian expression | 7.44E-04 |
| PGC-1a regulation | 8.22E-04 |
| Neuronal system | 8.22E-04 |
| Interleukin-2 signaling pathway | 8.22E-04 |
| NFAT involvement in hypertrophy of the heart | 8.22E-04 |
| Lipid metabolism regulation by peroxisome proliferator-activated receptor alpha (PPAR-alpha) | 8.22E-04 |
| Fatty acid beta oxidation | 0.001214738 |
| Cardiac muscle contraction | 0.001214738 |
| MAPK signaling pathway | 0.001309444 |
| Integrin signaling pathway | 0.001309444 |
| Mitochondrial beta-oxidation of saturated fatty acids | 0.001563155 |
| Calcium signaling pathway | 0.001563155 |
| Beta-1 integrin cell surface interactions | 0.001563155 |
| Semaphorin interactions | 0.001563155 |
| Mitochondrial fatty acid beta-oxidation | 0.001563155 |
| Valine, leucine and isoleucine degradation | 0.001954415 |
| EGFR interaction with phospholipase C-gamma | 0.002557237 |
| Phospholipase C-mediated cascade | 0.00300175 |
| Granule cell survival pathway | 0.003682381 |
| YAP1- and WWTR1 (TAZ)-stimulated gene expression | 0.003682381 |
| PAC1 receptor pathway | 0.00369064 |
| PDGFB signaling pathway | 0.004011975 |
| LPA receptor mediated events | 0.0043239 |
| RORA activates circadian expression | 0.004397835 |
| VEGFR1 pathway | 0.004397835 |
| Signal-dependent regulation of myogenesis by corepressor MITR | 0.004533691 |
| S1P/S1P1 pathway | 0.00475553 |
| MicroRNAs in cardiomyocyte hypertrophy | 0.005464493 |
| G-protein signaling pathways | 0.005870468 |
| Myometrial relaxation and contraction pathways | 0.005978282 |
| Neurotrophin signaling pathway | 0.006715603 |
| Heart development | 0.006715603 |
| Pathways in cancer | 0.006715603 |
| Signaling by VEGF | 0.006715603 |
| ALK in cardiac myocytes | 0.007418239 |
| Signaling by bone morphogenetic protein (BMP) | 0.007535325 |
| Signaling by FGFR in disease | 0.007574762 |
| Gap junction pathway | 0.008592747 |
| Alpha-6 beta-1 and alpha-6 beta-4 integrin signaling | 0.008663752 |
| Creatine metabolism | 0.008663752 |
| Myocyte adrenergic pathway | 0.008763638 |
| Calcineurin-dependent NFAT signaling role in lymphocytes | 0.009013836 |
| TAp63 pathway | 0.009013836 |
| BMP signaling and regulation | 0.009013836 |
| Hormone-sensitive lipase (HSL)-mediated triacylglycerol hydrolysis | 0.009013836 |
| Epidermal growth factor receptor (EGFR) pathway | 0.009013836 |
| Regulation of transcription by NOTCH1 intracellular domain | 0.009253901 |
| Signaling by PDGF | 0.009498475 |
| Signaling by ERBB2 | 0.009498475 |
| Insulin signaling pathway | 0.009744546 |
| Wnt signaling pathway | 0.009744546 |
| Differentiation pathway in PC12 cells | 0.010615589 |
| SHP2 signaling | 0.010982211 |
| Phosphatidylinositol signaling system | 0.015869133 |
| Prolactin regulation of apoptosis | 0.016009154 |
| BAD phosphorylation regulation | 0.016009154 |
| PPAR signaling pathway | 0.016009154 |
| Insulin secretion regulation by glucagon-like peptide-1 | 0.016769627 |
| Porphyrin metabolism | 0.016769627 |
| Sema3A-plexin repulsion signaling by inhibiting integrin adhesion | 0.016769627 |
| Signaling events mediated by VEGFR1 and VEGFR2 | 0.017340711 |
| EGFR1 pathway | 0.017881707 |
| G-protein-mediated events | 0.0188808 |
| Prion pathway | 0.019212146 |
| Signaling by EGFR in cancer | 0.019212146 |
| Ketone body biosynthesis and degradation | 0.019212146 |
| PPAR-gamma coactivator role in obesity and thermogenesis | 0.019212146 |
| Type I hemidesmosome assembly | 0.019212146 |
| Transmission across chemical synapses | 0.0232682 |
| MAPK cascade role in angiogenesis | 0.0240623 |
| BDNF signaling pathway | 0.024187351 |
| Gastric acid secretion | 0.024954057 |
| ECM-receptor interaction | 0.024954057 |
| Butanoate metabolism | 0.024954057 |
| MicroRNAs in muscle cell differentiation | 0.024954057 |
| Beta-oxidation of lauroyl-CoA to decanoyl-CoA | 0.024954057 |
| Ketone body metabolism | 0.024954057 |
| Neurophilin interactions with VEGF and VEGF receptor | 0.024954057 |
| PKA-mediated phosphorylation of key metabolic factors | 0.024954057 |
| Fibroblast growth factor receptor pathway | 0.024954057 |
| Agrin in postsynaptic differentiation | 0.024954057 |
| LKB1 signaling events | 0.024954057 |
| Interleukin-10 anti-inflammatory signaling pathway | 0.025098474 |
| LPA4-mediated signaling events | 0.025098474 |
| Sonic Hedgehog (Shh) pathway | 0.025098474 |
| Proximal tubule bicarbonate reclamation | 0.025447829 |
| Diurnally regulated genes with circadian orthologs | 0.027681865 |
| Adipocytokine signaling pathway | 0.028795752 |
| SIDS susceptibility pathways | 0.028795752 |
| TSH regulation of gene expression | 0.028795752 |
| Plexin D1 signaling | 0.030709001 |
| ATF2 transcription factor network | 0.0339296 |
| Potassium channels | 0.0339296 |
| Type 2 diabetes mellitus | 0.034502161 |
| Salivary secretion | 0.034502161 |
| Integrin beta-4 pathway | 0.034502161 |
| PIP2 hydrolysis | 0.036334209 |
| Ephrin receptor B forward pathway | 0.036625018 |
| Adipogenesis | 0.037208806 |
| BMP receptor signaling | 0.038045693 |
| PKA activation in glucagon signaling | 0.038478144 |
| N-cadherin signaling events | 0.041534013 |
| Signaling events mediated by stem cell factor receptor (c-Kit) | 0.041907808 |
| RXR/VDR pathway | 0.041907808 |
| TGF-beta signaling in development | 0.041907808 |
| Signaling events mediated by focal adhesion kinase | 0.043700842 |
| Nucleotide G-protein coupled receptors (GPCRs) | 0.044949093 |
| EGF receptor (ErbB1) signaling pathway | 0.046240427 |
| Interleukin-7 signaling pathway | 0.046240427 |
| Interleukin-4 signaling pathway | 0.046240427 |
| Signaling by NOTCH1 | 0.047422426 |
| ERBB signaling pathway | 0.04764733 |
| Mammalian calpain regulation of cell motility | 0.04764733 |
| Oxidative stress | 0.04764733 |
| Vascular smooth muscle contraction | 0.048870503 |
| Small cell lung cancer | 0.049028163 |
| Kit receptor signaling pathway | 0.049028163 |
| Integrins in angiogenesis | 0.049512105 |
| Wnt/calcium/cyclic GMP pathway | 0.049512105 |
| Angiotensin II-stimulated signaling through G-proteins and beta-arrestin | 0.049512105 |

## IRE1**α** inactivation led to loss of LV-specificity in vivo

To demonstrate the role of IRE1α as an upstream regulator of sXbp1 in neonatal heart, we randomly administered 4µ8C at 10mg/kg daily at P1 to P3 (early postnatal period) to wildtype neonatal mice when the robust differences in myocyte proliferation, growth, and programmed cell deaths were expected (**Figure 2A**). 4µ8C treatment significantly reduced sXbp1 in the heart and had no effect on total body weight (**Figure 2B and 2C**). Consistent with the differential expression pattern of sXbp1 between LV and RV (**Figure 1A-C**), 4µ8C specifically reduced LV weight but not RV weight (**Figure 2D**). To further characterize the effect of IRE1α inhibition on chamber-specificity, we performed proliferation, hypertrophic growth, and cell death assays by phospho-histone H3, WGA, and TUNEL staining, respectively. Consistent with previous report ^30^, the neonatal LV showed higher cardiomyocyte proliferation and hypertrophic growth than the RV, and lower cardiomyocyte death than the RV in the vehicle-treated hearts. In contrast, we observed reduced proliferation and hypertrophic growth in the 4µ8C-treated group in a LV-specific manner (**Figure 2E and 2F**). Meanwhile, 4µ8C-treatment increased cardiomyocyte death in the LV but not in the RV (**Figure 2G**).

**Figure 2.**
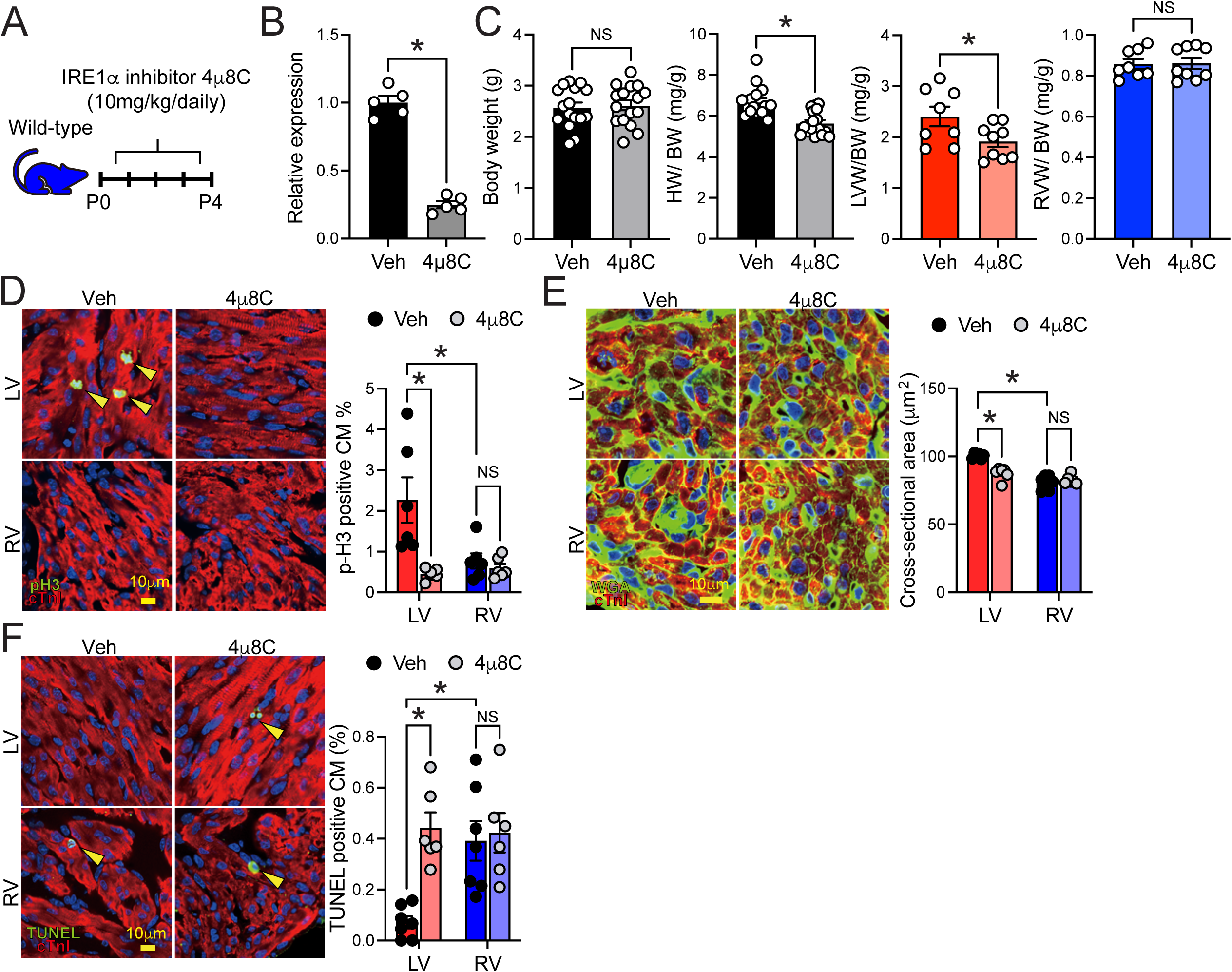
**Neonatal IRE1**α**-inactivated mice exhibit LV-specific dysregulation in chamber-specific behaviors. (A)** Experimental scheme of 4µ8C, an IRE1α inhibitor, administration to early postnatal (P1-P3) mice. **(B)** Down-regulation of sXbp1 in hearts from 4µ8C-treated mice at P4 (n=5). **(C)** Body weight (BW) and whole heart weight (HW) (n=16/Vehicle (Veh) and n=17/4µ8C) **(D)** LV weight (LVW) and RV weight (RVW)(n=8/Veh and n=9/4µ8C) **(E)** Immunohistochemistry against phospho-Histone H3, Wheat Germ Agglutinin (WGA) **(F)**, and TUNEL **(G)** (n=6). All data are shown as mean ± SEM, NS indicates not significant. *p<0.05

To demonstrate the temporal effect of IRE1α inhibition, we administered 4µ8C from P4 to P6 (mid-postnatal period), when ventricular growth shifts from proliferation to hypertrophy with minimal apoptosis (**Figure S2A**). Similar to the early postnatal inhibition, inhibition of IRE1α in P4-P6 neonatal heart also reduced LV weight specifically, but the change was more modest than that of early postnatal inhibition, with no significant changes in whole-heart weight detected (**Figure S2B and S2C**). Additional immunostaining also demonstrated that IRE1α inhibition at P4-P6 period also reduced cardiomyocyte proliferation and hypertrophic growth, and induced cardiomyocyte death specifically in the LV (**Figure S2D-F**). These data suggest that the IRE1α-sXbp1 axis regulates LV-specific cardiomyocyte growth during the first week of the perinatal period.

### Cardiac-specific IRE1**α** activation blunted chamber-specificity of ventricles and caused abnormal heart enlargement

Next, to further demonstrate the role of IRE1α in cardiomyocytes, we generated a cardiac-specific IRE1α transgenic mouse (IRE1α-cTg) by crossing Cre-dependent inducible IRE1α transgenic mice as reported previously ^33^ with the Nkx2.5-Cre mice ^34^ (**Figure S3A**). The animal exhibited marked induction of IRE1α and sXbp1 in the transgenic heart (**Figure S3B**). A robust enlargement of the heart was observed in the IRE1α-cTg at P7 and P28 time points(**Figure S3C and S3D**) withincreased size in both LV and RV (**Figure S3E**). Histological analysis revealed prominent chamber dilation (**Figure S3F**). To further determine the dynamic change of chamber morphology during the postnatal period, we performed serial echocardiograms at P1, P7, and P28. The IRE1α-cTg showed slight LV dysfunction (Control: 72.1±2.88 vs. IRE1α-cTg: 59.2±2.36) and dilation at P1, which progressed significantly at P7 (Control: 80.1±2.13 vs. IRE1α-cTg: 21.3±4.48) and later (**Figure S3G and S3H**). Immunohistochemical analyses revealed a robust induction of cardiomyocyte proliferation in both ventricles at P1 and P7 in the IRE1α-cTg in comparison to the controls (**Figure S3I**). Interestingly, the IRE1α-cTg mouse exhibited higher proliferative activity in the LV even at a late time point of P28 than the control group. (**Figure S3J**). In contrast, IRE1α-cTg blunted cardiomyocyte death in the RV at P1, an RV-specific feature during normal growth (**Figure S3K**). These data further demonstrate that IRE1α governs chamber-specific biological features in neonatal heart.

### Cardiac-specific Xbp1 deletion altered chamber-specificity of left ventricle

To further determine the role of Xbp1 in chamber-specific growth *in vivo*, we crossed the Myl2 (Mlc2v)-cre mice ^35^ with Xbp1 floxed mice ^36^ (cardiomyocyte-specific knockout [cKO]) (**Figure 3A**). The Xbp1-cKO neonates exhibited LV-specific weight reduction at P3, consistent with results observed with IRE1α inhibitor treatment (**Figure 3B and 3C**). As expected, Xbp1-cKO heart showed reduced proliferation and hypertrophic growth at P3 in a LV-specific manner (**Figure 3D and 3E**). In addition, Xbp1-cKO increased cardiomyocyte death in the LV but not in the RV (**Figure 3F**). These results further support that the IRE1α-Xbp1 axis regulates LV-specific myocyte growth.

**Figure 3.**
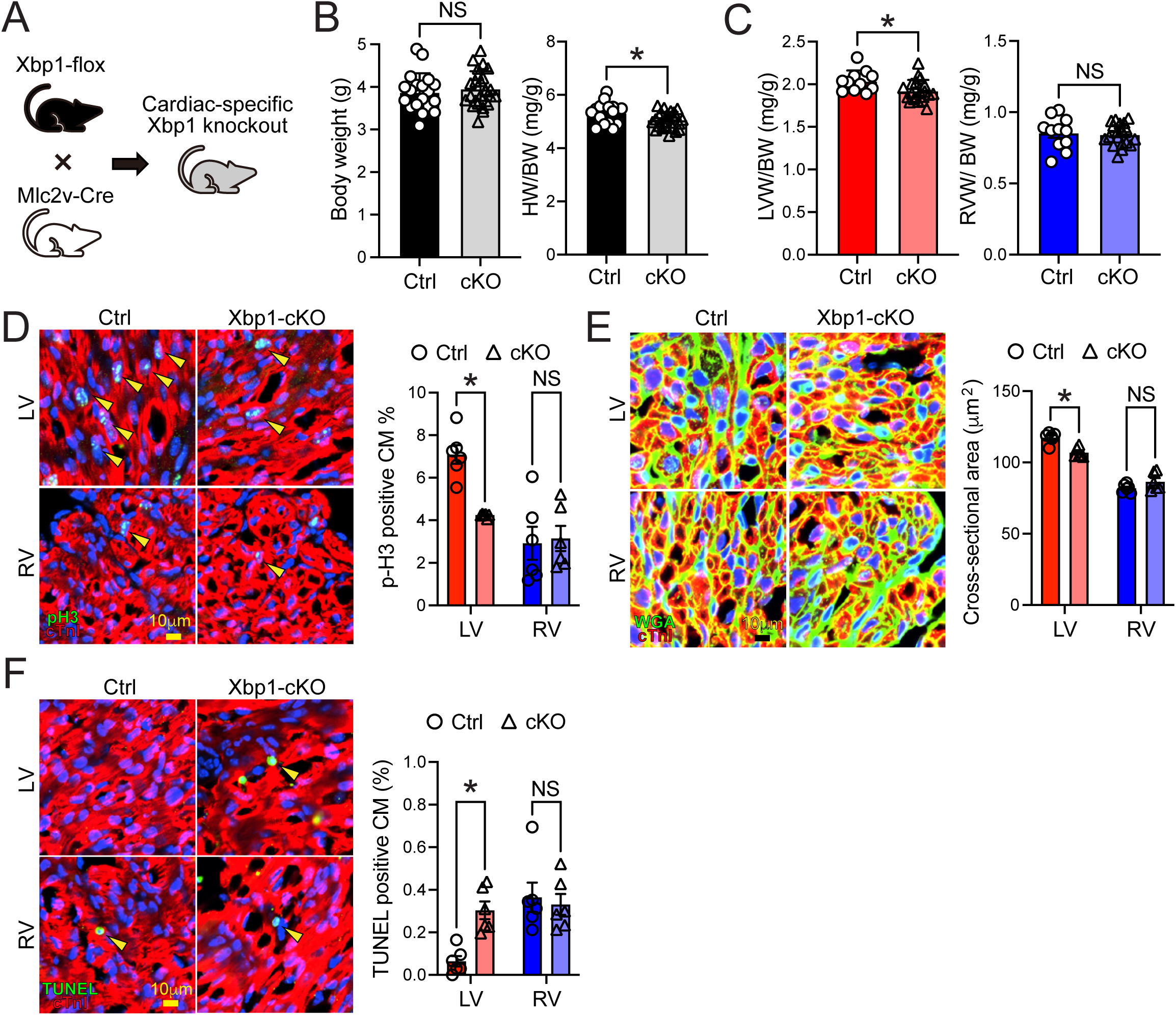
Neonatal Xbp1-inactivated mice exhibit LV-specific dysregulation in chamber-specific behaviors. **(A)** Breeding scheme of cardiac-specific Xbp1 knockout mice (Xbp1-cKO). **(B)** Body weight (BW) and whole heart weight (HW) (n=19/Ctrl and n=25/cKO) **(C)**, LV weight (LVW) and RV weight (RVW) at P7. (n=11/Ctrl and n=19/cKO) **(D-F)** Immunohistochemistry against phospho-Histone H3 (p-H3) **(D)**, Wheat Germ Agglutinin (WGA) **(E)**, and TUNEL **(F)** at P3 (n=6/Ctrl and n=6/cKO). All data are shown as mean ± SEM, NS indicates not significant, *p<0.05

### Cardiac-specific Xbp1 activation disrupted chamber-specificity of ventricles

To further elucidate the role of sXbp1 in the chamber-specific postnatal heart growth, we generated a cardiac-specific sXbp1 transgenic mouse (sXbp1-cTg) by crossing sXbp1-TRE ^37^ with Myh6-tTA ^38^ (**Figure 4A**). The sXbp1-cTg successfully induced sXbp1 in both ventricles at P3 (**Figure 4B**). The sXbp1-cTg showed increased whole heart, LV, and RV weights at P28 but not at P3 (**Figure 4C**), similar to the IRE1α-cTg hearts. Immunohistological analysis demonstrated increased proliferation but no change in hypertrophic growth in the sXbp1-cTg at P3 (**Figure 4D and 4E**), consistent with the observations made in the IRE1α-cTg hearts. In addition, sXbp1-cTg reduced cardiomyocyte death in the RV in the same manner as shown in the IRE1α-cTg (**Figure 4F**). Since hypertrophic growth is prominent at P7 and later timepoint, we conducted more histological analysis in the sXbp1-cTg at P28. sXbp1-cTg showed significant increases in cardiomyocyte sizes at P28 (**Figure 4G**). Interestingly, the sXbp1-cTg heart retained proliferative activity even at P28, which was also observed in the IRE1α-cTg hearts. Taken together, these in vivo results indicate that IRE1α and Xbp1 signal axis regulates differential growth between LV and RV during the postnatal period.

**Figure 4.**
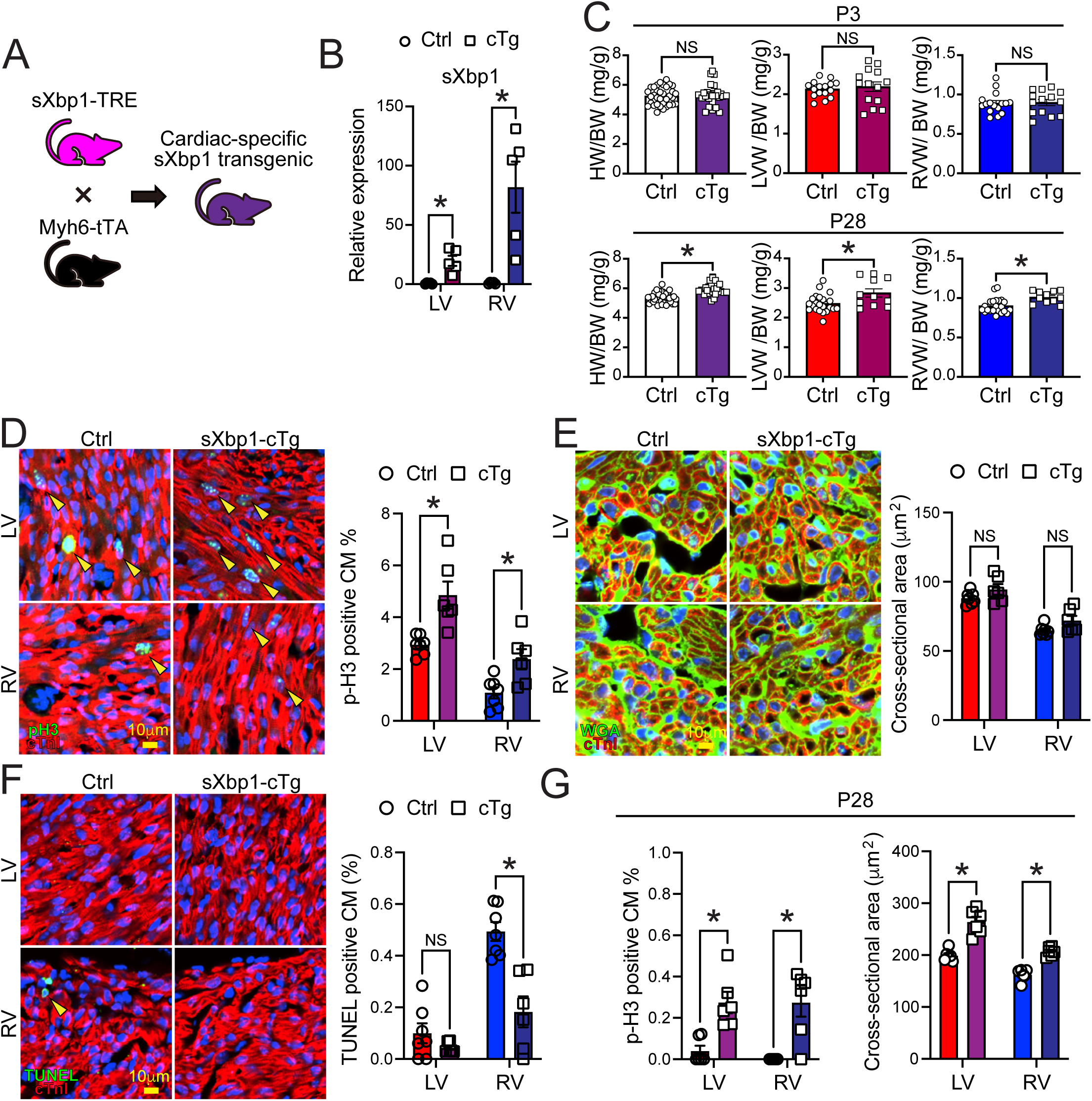
sXbp1 activation in cardiomyocytes leads to blunted chamber-specificity. **(A)** Breeding scheme of cardiac-specific Xbp1 induction mice (sXbp1-cTg). **(B)** sXbp1 expression in sXbp1-cTg and control (Ctrl) hearts at P3 (n=5). **(C)** Whole heart weight (HW), LV weight (LVW), and RV weight (RVW) at P3 and P28. (n=41/Ctrl and n=32/cTg for HW at P3; n=23/Ctrl and n=20/cTg for LVW and RVW at P3; n=36/Ctrl and n=23/cTg for HW at P28; n=22/Ctrl and n=11/cTg for LVW and RVW at P28) **(D-F)** Immunohistochemistry against phospho-Histone H3 (p-H3) **(D)**, Wheat Germ Agglutinin (WGA) **(E)**, and TUNEL **(F)** at P3 (n=7/Ctrl and n=6/cTg). **(G)** Quantification of immunohistochemistry against phospho-Histone H3 (p-H3) and Wheat Germ Agglutinin (WGA) at P28 (n=6/Ctrl and n=6/cTg). All data are shown as mean ± SEM, NS indicates not significant, *p<0.05

### Vimp and Rpn2 regulate sXbp1-dependent cardiomyocyte proliferation and hypertrophic growth

To uncover the underlying mechanisms for IRE1α-Xbp1 axis, we explored the downstream targets of Xbp1. sXbp1 specific ChIP-Seq was performed in the sXbp1-expressing neonatal cardiomyocytes *in vitro*. We obtained approximately 3,000 peaks, of which 67 were located in the gene promoter region. Among the 67 genes, 33 genes were up/downregulated in sXbp1-induced cardiomyocyte as determined by bulk RNA-seq (**Figure 5A**). From a published single-cell RNA-Seq data set ^39^ and we found that Xbp1 was abundant in the CM1 and CM2 subclusters associated with higher proliferative activity and less in the CM5 characterized as mature cardiomyocytes (**Figure 5B and C**). Among the 33 genes, we selected 6 top candidates, Rpn2, Vimp, Lrrc59, Dkc1, Slc30a5, and Odc1, based on their highly concordant gene expression pattern as Xbp1 in the subclusters of neonatal myocytes (**Figure 5D and S4A**). We validated the enrichment sXbp1 on the promoter regions of these candidate genes by ChIP-PCR in cultured neonatal cardiomyocytes expressing sXbp. VCP-Interacting Membrane Protein (Vimp; also known as Selenos) and Ribophorin II (Rpn2) were further characterized since their expression was detected at high levels in the proliferating cardiomyocyte subpopulation based on single cell transcriptome data (CM-1,-2, and-4)( **Figure 5E and S4B**). In addition, we analyzed the gene expression profiles in the Rpn2-or the Vimp-expressing cardiomyocytes from the scRNA-Seq dataset and found that both genes were positively related to proliferation and negatively related to maturation in postnatal cardiomyocytes (**Figure S4C**). To determine direct interaction of Xbp1 with these two genes *in vivo,* we performed Xbp1-ChIP using isolated cardiomyocytes from sXbp1-cTg hearts at P3 and detected significant enrichment of Xbp1 interaction with both Vimp and Rpn2 promoters (**Figure 5F**).

**Figure 5.**
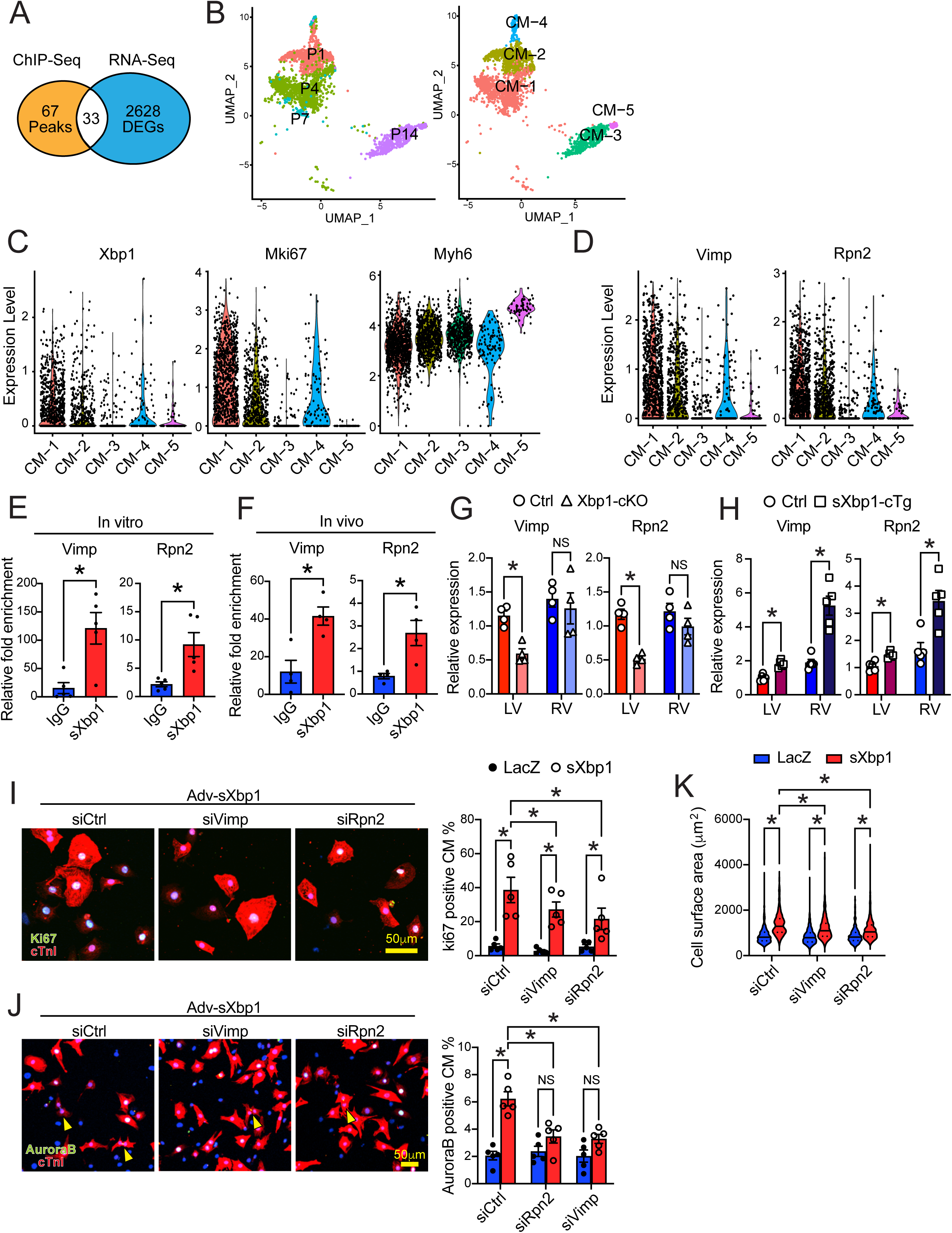
Identification of sXbp1 downstream targets, Vimp and Rpn2, and their role in neonatal cardiomyocytes. **(A)** Venn diagram from sXbp1-chromatin immunoprecipitation (ChIP) followed by sequencing and bulk RNA-Seq datasets in sXbp1-induced neonatal cardiomyocytes. **(B)** Subclustering of cardiomyocyte population from mouse heart at P1, P4, P7, and P14. **(C)** Violin plots for Xbp1, Mki67 (proliferation marker), and Myh6 (maturation marker) in cardiomyocyte sub-populations. **(D)** Violin plots for Vimp and Rpn2 in cardiomyocyte sub-populations. **(E, F)** sXbp1-ChIP followed by qPCR against Vimp and Rpn2 in sXbp1-induced neonatal cardiomyocytes **(E)** (n=5) and in cardiomyocytes isolated from cardiac-specific sXbp1-induced hearts **(F)** (n=4). **(G,H)** Vimp and Rpn2 expression in cardiac-specific Xbp1 knockout (Xbp1-cKO) **(G)** and cardiac-specific sXbp1-induced (sXbp1-cTg) hearts **(H)**. **(H)** Contraction forces generated by either Control or Col5a1CKO CFs (n=3) (**I-K**) Determination of contractile forces by generating Col5a1CKO CFs ex vivo. **(I,J)** Immunocytochemistry against Ki67 (I) and Aurora B (J) in neonatal cardiomyocytes treated with adv-sXbp1 in the presence of siRNA against Vimp and Rpn2 (n=5). **(K)** Cell surface area of neonatal cardiomyocytes treated with adv-sXbp1 and siRNAs (n=5). All data are shown as mean ± SEM, NS indicates not significant, *p<0.05

Furthermore, both Vimp and Rpn2 were downregulated specifically in the LV in both Xbp1-cKO and IRE1α-cKO hearts (**Figure 5G and S4D**), while significantly enhanced in both ventricles in the IRE1α and Xbp1 expressing hearts (**Figure 5H and S4E**). All these results support the notion that Vimp and Rpn2 are downstream targets of IRE1α-Xbp1 axis in neonatal myocytes.

Both Vimp and Rpn2 are endoplasmic reticulum (ER) targeted transmembrane proteins. In beta cells and adipocytes, Vimp is known to regulate proliferation and survival^40,41^. Rpn2 has emerged as an important regulator of cell proliferation and survival as well as tumor growth and metastasis in cancers^42–44^. To demonstrate the role of Vimp and Rpn2 in cardiomyocyte *in vitro*, we treated neonatal cardiomyocytes with siRNA against Vimp or Rpn2 in combination with sXbp1 over-expression (**Figure S4F**) and observed that sXbp1-induced proliferation was significantly blunted by gene silencing of Rpn2 or Vimp (**Figure 5I and 5J**). In addition, sXbp1-induced hypertrophic growth was also blunted by the gene silencing of Vimp or Rpn2 (**Figure 5K**). These results clearly demonstrated that Rpn2 and Vimp are essential downstream mediators for sXbp1-induced proliferation and hypertrophic growth of neonatal cardiomyocytes.

### *In vivo* function of Vimp or Rpn2 in neonatal heart growth

To validate the cell-autonomous role of Rpn2 and Vimp *in vivo*, we generated mosaic mutants of each gene using CRISPR/Cas9/AAV9-based somatic mutagenesis ^45^ (**Figure 6A and S5A**). To achieve approximately 50% AAV transduction rate, we injected 2.0 x 10^10^ vg/g of AAV9-cTnT-gRNA-Vimp (AAV9-sgVimp) or AAV9-cTnT-gRNA-Rpn2 (AAV9-sgRpn2) into Rosa26^Cas9-GFP^ P1 pups ^46^ and harvested the heart at P7. Using this approach, we successfully achieved 39.4% GFP-positive cardiomyocytes in the AAV9-sgVimp-administered group and 49.3% in the AAV9-sgRpn2-administered group (**Figure 6B and 6C**). To determine the impact of Vimp and Rpn2 deletions on proliferation, we stained AAV9-injected Rosa26cas9-GFP hearts with phospho-histone H3 (p-H3) and checked p-H3-positive cardiomyocytes between control (GFP-negative) and mutant (GFP-positive) cardiomyocytes. AAV9-sgVimp-injected Rosa26^Cas9-GFP^ (Vimp-Cas9) hearts showed a significant decrease in p-H3-positive cells in mutant cardiomyocytes compared with the control in the LV (Control: 4.98±0.37% vs. Mutant: 2.33±0.44%) (**Figure 6D**). We also noted a decrease in proliferation within the RV (Control: 2.46±0.20% vs. Mutant: 1.40±0.28%). Additionally, hearts injected with AAV9-sgRpn2 in Rosa26Cas9-GFP (Rpn2-Cas9) also displayed a significant reduction in proliferation of mutant cardiomyocytes compared to controls in the LV (Control: 4.65±0.33% vs. Mutant: 2.11±0.16%) (**Figure 6E**). Again, we also observed a reduction in proliferation in the RV (Control: 2.24±0.13% vs. Mutant: 1.33±0.23%), similar to the Vimp-chimeric mutant hearts. WGA staining showed that both Vimp-Cas9 and Rpn2-Cas9 hearts had decreased cardiomyocyte sizesspecifically in the LV of the mutants (Control: 117±2.33% vs. Mutant: 96.7±2.25% in Vimp-Cas9; Control: 119±3.21% vs.

**Figure 6.**
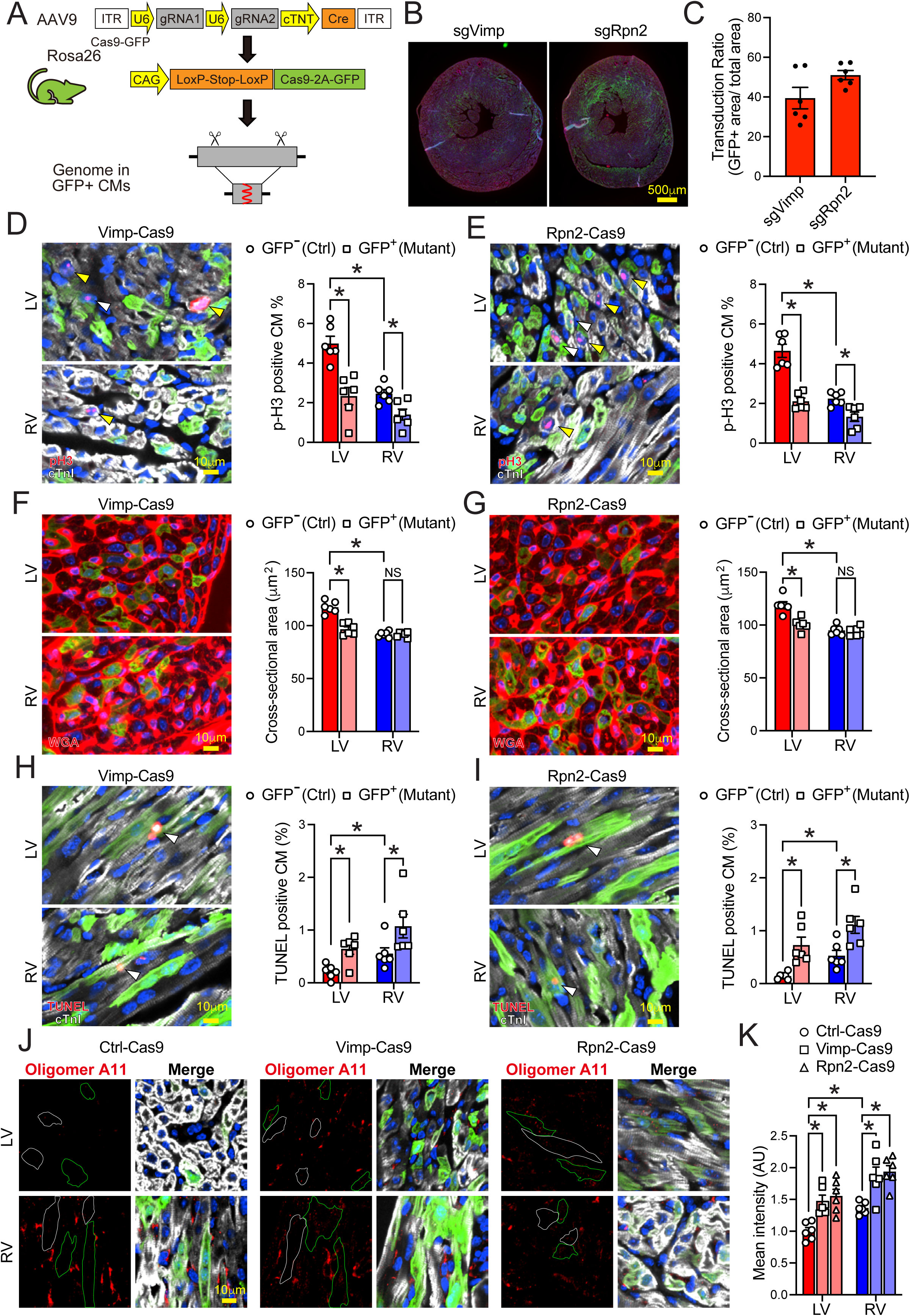
Mosaic mutagenesis mouse models against Vimp and Rpn2. **(A)** Generation of mosaic mutants using CRISPR/Cas9/AAV9-based somatic mutagenesis. (**B,C**) Representative images of mosaic mutant hearts **(B)** and transduction ratio **(C)** (n=6). **(D,E)** Immunostaining for phospho-Histone H3 (p-H3) in Vimp (Vimp-Cas9) **(D)** and Rpn2 (Rpn2-Cas9) **(E)** mutant hearts. (Yellow arrows indicate p-H3-positive “control” [GFP-negative] cardiomyocyte, and white arrows indicate pH3-positive “mutant” [GFP-positive] cardiomyocyte) (n=6). **(F,G)** Immunostaining for Wheat Germ Agglutinin (WGA) in Vimp-Cas9 **(F)** and Rpn2-Cas9 **(G)** mutant hearts (n=6). **(H,I)** Immunostaining for TUNEL in Vimp-Cas9 **(H)** and Rpn2-Cas9 **(I)** mutant hearts (White arrows indicate TUNEL-positive “mutant” [GFP-positive] cardiomyocyte) (n=6). **(J,K)** Representative images of Oligomer A11 **(J)** and quantification **(K)** of mosaic mutant hearts (n=6). Control cardiomyocytes are highlighted in white and mutant cardiomyocytes are highlighted in green. All data are shown as mean ± SEM, NS indicates not significant, *p<0.05

Mutant: 101±2.43% in Rpn2-Cas9) (**Figure 6F and 6G**). Furthermore, both Vimp-Cas9 and Rpn2-Cas9 hearts exhibited increased cell death in mutant cardiomyocytes within the LV (Control: 0.21±0.05% versus Mutant: 0.64±0.10% in Vimp-Cas9; Control: 0.14±0.03% vs. Mutant: 0.73±0.15% in Rpn2-Cas9), as well as in the RV (Control: 0.55±0.11% vs. Mutant: 1.07±0.22% in Vimp-Cas9; Control: 0.53±0.10% vs. Mutant: 1.11±0.16% in Rpn2-Cas9) (**Figure 6H and 6I)**.

Since both Vimp and Rpn2 are associated with ER protein homeostasis ^47–51^, we evaluated aggregated protein levels in the mutant hearts by detecting preamyloid oligomers (PAOs). We found that both mutants showed increased PAOs (**Figure 6J and 6K**). Furthermore, the RV showed a higher PAO accumulation than the LV in the control hearts.

These results suggest that Vimp and Rpn2 are two downstream targets of Xbp1 that regulate differential growth between the RV and the LV during the postnatal period, potentially involving protein homeostasis.

## Discussion

The transition from fetal to postnatal life is characterized by dramatic shifts in hemodynamic loading, prompting the left ventricle (LV) and right ventricle (RV) to undergo divergent morphological and functional changes^1–8^. While the role of extrinsic mechanical forces in driving this chamber-specific remodeling is well-recognized, the intrinsic intracellular networks that orchestrate these precise asymmetric growth patterns have remained elusive^9–12^. In this study, we identify the endoplasmic reticulum (ER) stress-responsive IRE1α-Xbp1 signaling pathway as a fundamental, intrinsic determinant of LV-specific postnatal development. We demonstrate that the IRE1α-sXbp1-Vimp/Rpn2 axis specifically drives the elevated cardiomyocyte proliferation, hypertrophic growth, and enhanced cell survival required for the rapid expansion of the LV after birth (**Figure S6A**).

### The Role of UPR in Chamber-Specific Postnatal Growth

Our findings reveal a previously underappreciated developmental role for the unfolded protein response (UPR) in the heart^19–22^. The LV’s sudden exposure to high systemic vascular resistance at birth demands a rapid, massive increase in protein synthesis to support cardiomyocyte hypertrophy and proliferation^3–6,13^. This translational surge inherently challenges ER folding capacity. Our data show that the IRE1α-Xbp1 pathway is highly activated in the LV immediately after birth, likely serving as a crucial adaptive mechanism to manage this increased protein load. When this pathway is disrupted— either through pharmacological inhibition of IRE1α or genetic ablation of Xbp1—the LV fails to achieve its normal mass, showing reduced proliferation and hypertrophic growth alongside increased apoptosis. Strikingly, these loss-of-function interventions do not significantly impact the RV, highlighting the LV’s unique dependency on robust ER stress signaling for its normal postnatal morphogenesis. Conversely, forced cardiac-specific activation of IRE1α or sXbp1 blunts the natural chamber specificity, driving the RV to adopt an LV-like growth phenotype characterized by aberrant enlargement, sustained proliferation, and suppressed apoptosis.

### Mechanistic Insights: The Vimp and Rpn2

To elucidate the downstream effectors of this UPR-driven morphogenic program, we identified VCP-Interacting Membrane Protein (Vimp, also known as Selenos) and Ribophorin II (Rpn2) as direct transcriptional targets of sXbp1^40–44^. Both proteins are localized to the ER and are deeply involved in protein homeostasis; Vimp is a key component of ER-associated degradation (ERAD) ^47–49^, while Rpn2 is part of the oligosaccharyltransferase (OST) complex involved in N-linked glycosylation ^50,51^ and a regulatory subunit of the 26S proteasome, which is essential for the ATP-dependent degradation of ubiquitinated proteins^52–54^.

Our single-cell data analysis and in vitro validations confirm that Vimp and Rpn2 expression is tightly correlated with highly proliferative, less mature cardiomyocyte subpopulations. Furthermore, in vivo CRISPR/Cas9 somatic mutagenesis demonstrated that localized deletion of either Vimp or Rpn2 severely impairs cardiomyocyte proliferation and hypertrophy while increasing cell death, confirming their essential roles in executing the sXbp1-mediated growth program. By upregulating Vimp and Rpn2, sXbp1 enhances the ER’s capacity to process and quality-control the massive influx of newly synthesized structural and functional proteins required for postnatal ventricular expansion in the LV (**Figure S6B)**.

### Clinical Implications for Single-Ventricle Physiology

The discovery of this chamber-specific growth axis has profound clinical implications, particularly for congenital heart defects characterized by single-ventricle physiology, such as Hypoplastic Left Heart Syndrome (HLHS). In these conditions, surgical palliation (e.g., the Norwood procedure followed by Glenn and Fontan operations) forces the RV to support the systemic circulation ^55,56^. Unfortunately, the systemic RV frequently fails over time, as it intrinsically lacks the hypertrophic and proliferative capacity of a normal LV ^57,58^.

Our data suggest that the “RV-ness” or “LV-ness” of a chamber is not rigidly fixed but is governed by manipulable intracellular signaling pathways. Modulating protein homeostasis through the targeted activation of the IRE1α-Xbp1-Vimp/Rpn2 axis could represent a novel therapeutic strategy. By safely upregulating this pathway in the systemic RV of congenital heart disease patients, it may be possible to induce adaptive, LV-like hypertrophic and proliferative growth, thereby improving long-term systemic ventricular function and survival.

### Limitations and Future Directions

While our study provides robust in vivo and in vitro evidence, several questions remain. The upstream triggers that specifically activate the IRE1α-Xbp1 pathway in the LV, whether they are purely mechanical (stretch/afterload) or neurohormonal, require further investigation. Additionally, while transient activation of this pathway promotes adaptive growth postnatally, chronic or excessive UPR activation is a known driver of pathological hypertrophy and heart failure in adults^24–29^. Future studies must define the optimal therapeutic window and magnitude of pathway modulation to safely promote adaptive remodeling without triggering pathological ER stress-induced apoptosis.

## Conclusion

In summary, we have defined the IRE1α-Xbp1-Vimp/Rpn2 axis as a critical, chamber-specific regulator of postnatal cardiac growth. By linking ER protein homeostasis to cellular proliferation, hypertrophy, and survival, this pathway answers a longstanding question regarding the molecular origins of LV and RV asymmetry, offering a promising new biological target for single-chamber heart diseases.

## Author Contributions

BZ performed experiments and analyzed data. JH analyzed bulk RNA-and scRNA-Seq data. DC analyzed ChIP-Seq data. MJ, IR, and ML assisted in bench experiments and animal maintenance. JL assisted data analysis on IHC and ICC. AA isolated neonatal rat ventricle cardiomyocytes. AA, TH, and YW provided key reagents and contributed to the discussion; TY conceptualized the project, performed experiments, supervised data collection and analysis, and wrote the manuscript.

## Acknowledgements

We thank Drs. Zhao Wang and Yingfeng Deng, City of Hope Medical Center, for providing us Adv-sXbp1, the sXbp1-TRE mice, and the Xbp1-flox mice.

## Methods

### Animal care and use

All animal studies were approved by the Animal Research Committee, University of California, Los Angeles. All animals were maintained at the UCLA vivarium according to the policies instituted by the American Association for Accreditation of Laboratory Animal Care. Both Male and female animals were used in the study. All animals belonged to the C57Bl/6 strain, were healthy, immune-free, and drug or test naïve and were not involved in other experimental procedures. Littermates were used as controls for all experiments. Detailed experimental procedures are included in the Supplemental Material.

### Statistical Analysis

All data are represented as mean ± standard error of the mean (SEM). The exact value of n is mentioned in the figure legends and always stands for separate biological replicates. The number of animals for each experiment was calculated based on a power analysis approach. To detect a 50% change in significance using a 5% (0.05) p-value for a parameter with 20-30% biological variation and 80% power, the minimum number is 3-6 per group. Statistical analysis was performed using GraphPad (Prism) software using Student’s t-test (Two tailed) and one-way ANOVA with Tukey’s multiple comparison analysis as appropriate. A P value <0.05 was considered statistically significant.

### Data and Code Availability

The accession numbers for the sXbp1 bulk RNA sequencing and sXbp1-ChIP sequencing in this paper are available in the NCBI BioProject (PRJNA1129185). All code used in the bioinformatics analysis can be provided on request.

## Non-standard Abbreviations and Acronyms

AAV9: Adeno-associated virus 9
Adv: Adenovirus
ATF6: Activating transcription factor 6
ChIP: Chromatin immunoprecipitation
cTnT: Cardiac Troponin T
Dkc1: Dyskerin Pseudouridine Synthase 1
ER: Endoplasmic reticulum
ERAD: ER-associated degradation
gRNA: Guide RNA
HLHS: Hypoplastic Left Heart Syndrome
IRE1a: Inositol-requiring enzyme 1α
Lrrc59: Leucine Rich Repeat Containing 59
LV: Left ventricle
MAPK: Mitogen-activated protein kinase
mTOR: Mechanistic target of rapamycin kinase
Myh6: Myosin heavy chain 6
Myl2 (Mlc2v): Myosin Light Chain 2
Nkx2.5: NK2 homeobox 5
Odc1: Ornithine Decarboxylase 1
OST: Oligosaccharyltransferase
PAOs: Preamyloid oligomers
PERK: Eukaryotic translation initiation factor 2 alpha kinase 3
RNA-FISH: RNA-fluorescence in situ hybridization
Rpn2: Ribophorin II
RV: Right ventricle
scRNA-Seq: Single cell RNA sequencing
Slc30a5: Solute Carrier Family 30 Member 5
TRE: Tetracycline-responsive promoter element
tTA: Tetracycline-controlled transactivator
TUNEL: Terminal deoxynucleotidyl transferase dUTP nick end labeling
UPR: Unfolded protein response
Vimp: VCP-Interacting Membrane Protein
WGA: Wheat germ agglutinin
XBP1: X-Box Binding Protein 1

